# Retinoic acid production via the ray-finned fish gene *beta-carotene oxygenase 1-like* is essential for juvenile development

**DOI:** 10.1101/2025.09.30.679646

**Authors:** Lochlan Sife Krupa, Paula R. Villamayor, Sepalika Bandara, Yanqi Zhang, Alec Palmiotti, Johannes von Lintig, Andres Bendesky

## Abstract

In vertebrates, vitamin A (VA) is crucial for development, tissue homeostasis, vision, and immunity. Retinal, a form of VA, is produced via enzymatic cleavage of β-carotene by *beta-carotene oxygenase 1* (*bco1*) and *bco1-like* (*bco1l*). While *bco1* is found across vertebrate taxa, *bco1l* is a paralog of *bco1* that we discover to have evolved in the ray-finned fishes, the most abundant, speciose, and commercially important group of fishes. We investigated the function of *bco1l* in ray-finned Siamese fighting fish, commonly known as betta, an emerging model for genetics and development. Using CRISPR-Cas9 knockouts, we find that lack of *bco1l* results in reduced VA and elevated β-carotene in larvae, starting when animals have exhausted their yolk supply of retinal, followed by stunted growth and death during juvenile development. Exogenous retinoic acid rescues the mutation, demonstrating its deficiency causes these defects. *bco1l* is 4× more abundant than *bco1* in the intestine. This, coupled with the inability of *bco1* to sustain VA production in the *bco1l* mutant, indicates that *bco1l* is the primary enzyme for dietary carotenoid conversion into retinal. Our results show that VA production by *bco1l* is required for post-embryonic development, and that *bco1l* became essential after evolving via duplication of *bco1*.

## Introduction

Vitamin A (VA) deficiency has detrimental consequences, including improper pattern formation during development, stunted growth, compromised immunity, and impaired vision including blindness (Clagett-Dame and DeLuca 2002; Hernandez and Hardy 2020; Huang et al. 2018; Menezes and Almeida 2024). To prevent such defects, VA must be obtained preformed from the diet or enzymatically produced through the conversion of dietary pro-VA carotenoids (O’Byrne and Blaner 2013).

Due to their biological activities, carotenoids have a variety of applications. They are commonly employed as food supplements in aquaculture, where they are used to enhance animal coloration and growth (Helgeland et al. 2019; Liao et al. 2025). Carotenoids also have applications in medicine including boosting immune function, reducing risk of chronic disease, and treating VA deficiency (Metibemu and Ogungbe 2022; Vílchez et al. 2011; von Lintig 2010)

In vertebrates, the enzyme beta-carotene oxygenase 1 (encoded by the *bco1* gene) catalyzes the cleavage of the carotenoid β-carotene into two molecules of retinal, a form of VA, via 15,15’ oxygenase activity (**Fig. 1**; Harrison & Kopec, 2020; Wyss et al., 2000). Retinal is then irreversibly oxidized to the biologically active metabolite retinoic acid (RA) or reduced to retinol. Retinol can be further converted into retinyl esters, which are storage forms of VA (O’Byrne and Blaner 2013; von Lintig 2010). Retinyl esters also serve as precursors of the visual chromophore (11-*cis*-retinal) (Kiser et al. 2014). *bco1* is expressed in the epithelium of the small intestinal mucosa, where it contributes to producing VA from dietary β-carotene (Lindqvist and Andersson 2002; Takitani et al. 2006; Goodman and Huang 1965; Olson and Hayaishi 1965). This intestinal-derived VA is typically stored in the liver as retinyl esters (Lindqvist and Andersson 2004; von Lintig et al. 2005). *bco1* is also expressed in other tissues including the kidneys, lungs, liver, and retina, where it likely provides VA locally (Lindqvist and Andersson 2004; Mora et al. 2004; Paik et al. 2001; Wyss et al. 2001; Yan et al. 2001). Knockout of *bco1* in mice results in the accumulation of β-carotene, a reduction of VA stores, and an exacerbation of defects resulting from VA deficiency during embryonic development (Hessel et al. 2007; Kim et al. 2011). Morpholino-mediated knockdown of *bco1* in zebrafish results in malformation of neural crest derivatives such as the mandible and pigment cells during embryonic development (Lampert et al., 2003). However, a CRISPR/Cas9-mediated loss-of-function mutation in *bco1* in the closely related pearl danio only leads to a subtle pigmentation phenotype in adults (Huang et al. 2021). Therefore, the function of *bco1* in fish might be particularly important in tissues derived from the neural crest.

**Figure 1.**
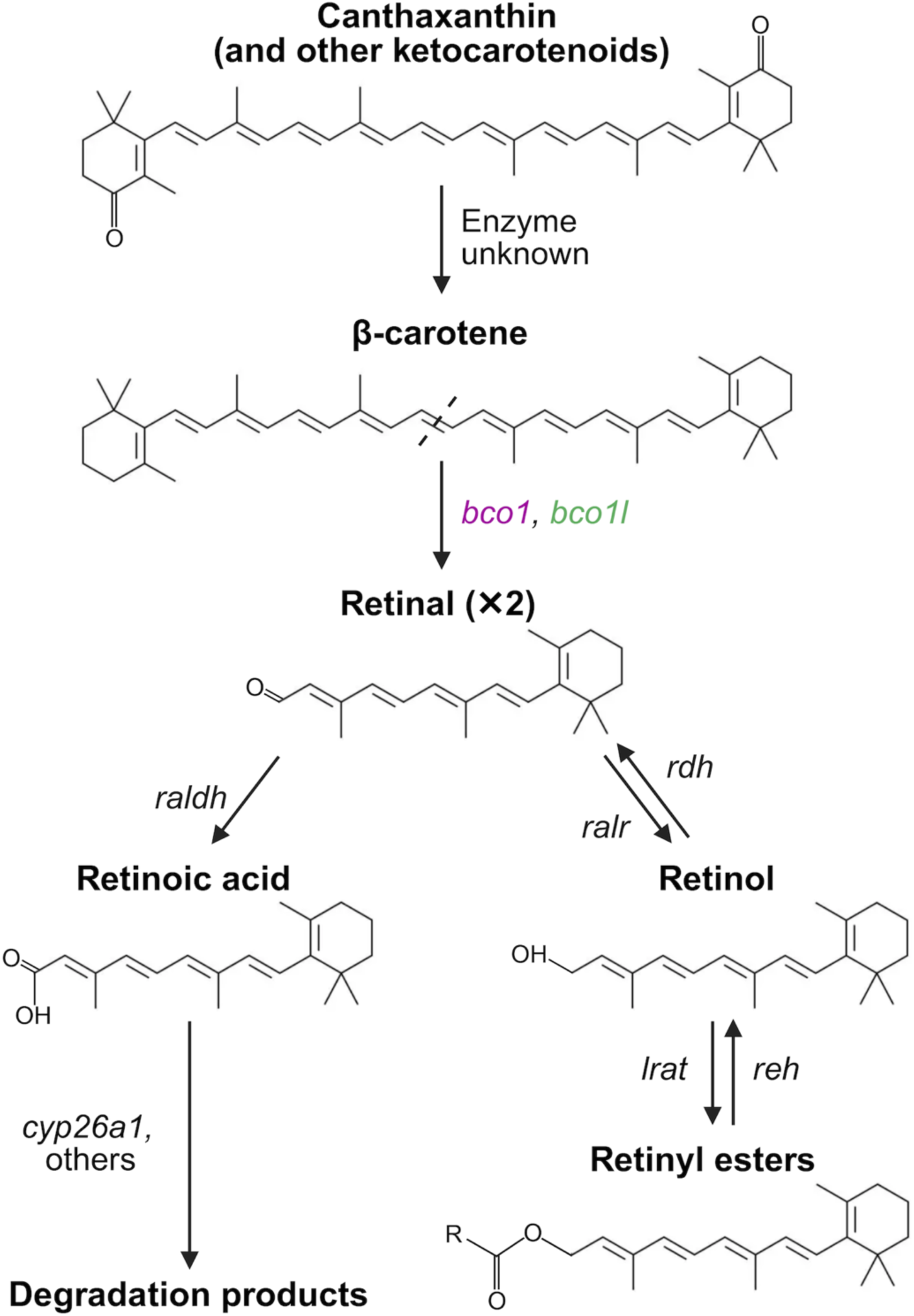
Metabolism of carotenoids to retinoids in vertebrates. Ketocarotenoids, such as canthaxanthin (pictured) and astaxanthin, can be converted to β-carotene in some species, though the enzymes involved in this process remain undescribed (Elshafey et al. 2023; Sangeetha and Baskaran 2010). Furthermore, there is evidence some species are incapable of this conversion. For example, astaxanthin lacks provitamin A activity in humans (Kumar et al. 2022), suggesting it is not readily converted to the provitamin A β-carotene. β-carotene is converted to two molecules of retinal via the 15,15’oxygenase activity of *bco1*. *bco1l* is also capable of catalyzing this reaction in the ray-finned fishes (Helgeland et al. 2019; Kwon et al. 2022). Retinal is converted to retinoic acid by one of three retinal dehydrogenases (*raldh*) (Lampert et al. 2003; O’Byrne & Blaner 2013). Retinoic acid is converted to inactive metabolites through the action of the cytochrome P450 enzymes such as *cyp26a1* (Isken et al. 2007; White et al. 1996). Retinal may also be reversibly reduced to retinol by retinal reductase (*ralr*), which is further acylated to the storage form (retinyl esters) by lecithin:retinol acyltransferase (*lrat*). Retinyl esters are converted to retinol by retinyl ester hydrolase (*reh*), which is oxidized to retinal by one of the retinol dehydrogenase (*rdh*) family members (O’Byrne and Blaner 2013).

*Beta-carotene oxygenase 1-like* (*bco1l*) is a paralog of *bco1* found in ray-finned fishes (*Actinopterygii*) (Helgeland et al. 2014). Like *bco1*, *bco1l* also catalyzes the symmetric cleavage of β-carotene to retinal (**Fig. 1**; Helgeland et al., 2019; Kwon et al., 2022). The presence of *bco1l* across ray-finned fishes is notable given that one of a pair of paralogous genes is typically pseudogenized following duplication (Lynch and Conery 2000). The preservation of both *bco1* and *bco1l* suggests that subfunctionalization (the division of the functions of the ancestral and derived paralogs) or neofunctionalization (the emergence of a novel function for one of the paralogs) may have occurred (Magadum et al. 2013). Several studies in Atlantic salmon provide preliminary evidence for the subfunctionalization of *bco1* and *bco1l*. The two genes display differential expression in the intestine, liver, and muscle (Helgeland et al. 2014) and may have distinct impacts on carotenoid deposition in muscle (Helgeland et al. 2019; Kuhn 2022). However, duplicate genes can be maintained for extended periods of time through processes such as dosage-balance selection and hypofunctionalization without undergoing subfunctionalization (Qian et al. 2010; Wilson and Liberles 2023), and the relative expression of paralogous genes can be subject to drift (Thompson et al. 2016). To date, functional roles and evolutionary trajectories of *bco1* and *bco1l* in ray-finned fish remains largely unknown.

We previously used CRISPR/Cas9 to knock out *bco1l* in betta fish, and observed that the homozygous mutant genotype was lethal (Palmiotti et al. 2023). In this study, we sought to characterize the developmental and biochemical phenotypes of the knockout and to parse the evolution and respective roles of *bco1* and *bco1l* in carotenoid metabolism and VA production.

## Results

### bco1l is required for juvenile but not embryonic or early larval development

While the consequences of a lack of *bco1* have been studied in zebrafish and mice (Hessel et al. 2007; Kim et al. 2011; Lampert et al. 2003), those of a 1 *bco1l* knockout have not been documented. Upon discovering that homozygosity of a *bco1l* knockout mutation was lethal in betta (Palmiotti et al. 2023), we set out to characterize the phenotypic consequences of the mutation and to identify when in development the mutants were dying. We initially imaged and genotyped embryos and larvae from a heterozygous × heterozygous cross at 3, 6, 10, 12, and 20 days postfertilization (dpf). Genotype ratios of the offspring did not differ significantly from the expected Mendelian ratios at any time point (**Fig. 2A**; Fisher’s exact tests, p > 0.05), indicating that there was no mortality associated with the knockout up to 20 dpf. Furthermore, we found no visible morphological defects associated with the knockout at any time point (**Supplemental Fig. S1**).

**Figure 2.**
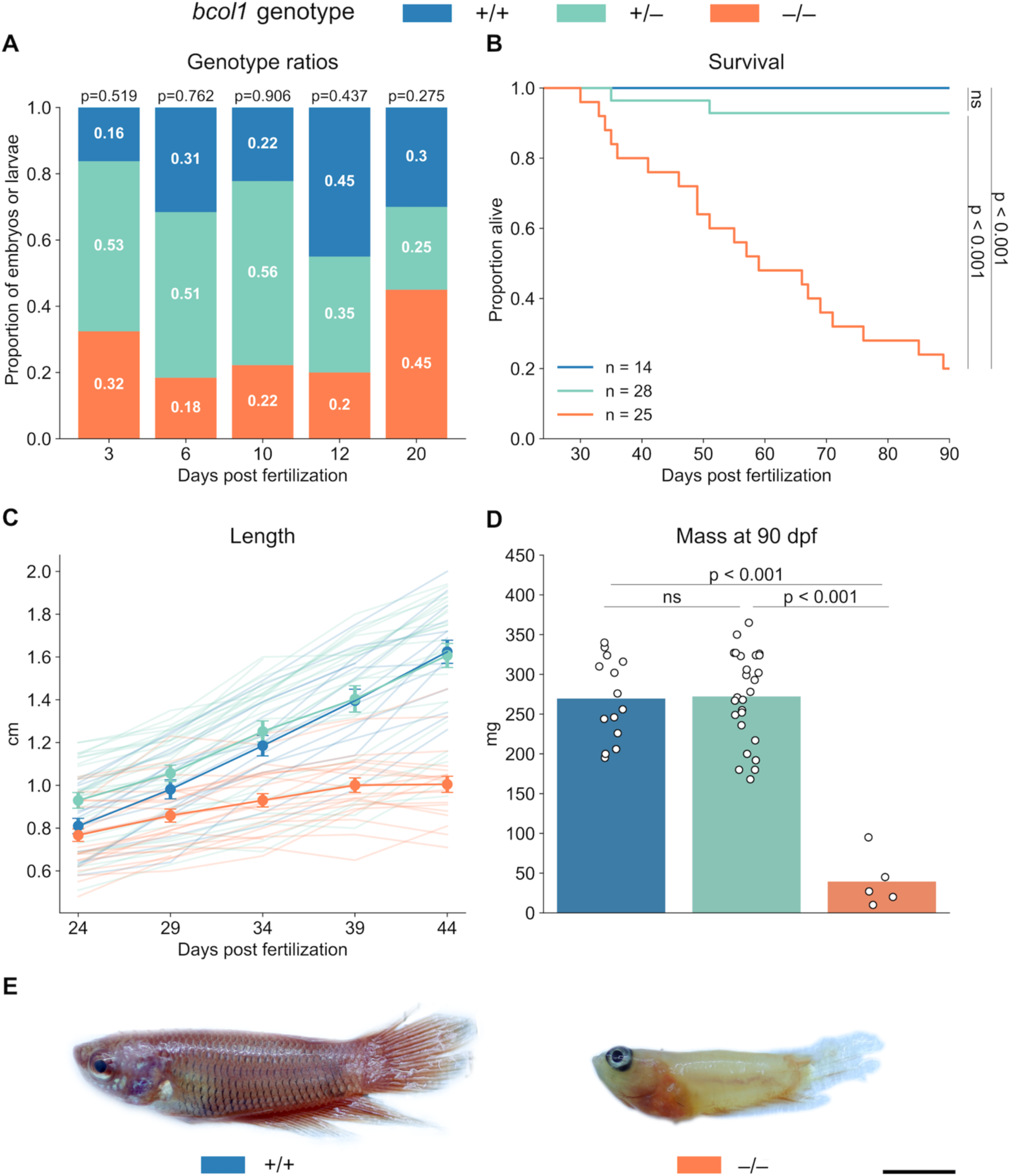
*bco1l* is required for survival and growth in betta fish. **A)** Genotype ratios of fish sampled at 3, 6, 10, 12, and 20 dpf from a heterozygous cross. There was no significant deviation from the expected Mendelian ratios at any time point. p-values by Fisher’s exact test. **B)** Survival curves for the three genotypes from 24 to 90 dpf. p-values by log-rank test adjusted by Bonferroni correction. **C)** Lengths of fish of the three genotypes from 24 to 44 dpf. Faint lines track the growth of individual fish; circles denote the mean ± s.e.m. **D)** Masses of surviving fish of the three genotypes at 90 dpf. p-values by one-way ANOVA with Tukey’s HSD. **E)** Representative images of +/+ and –/– individuals at 90 dpf. Scale bar: 5mm.

To determine at what age after 20 dpf the –/– fish were dying and to document their larval and juvenile development, we individually housed larval offspring from a heterozygous cross and measured growth and survival from 24 to 90 dpf. The –/– fish had impaired survival relative to both +/+ (**Fig. 2B**; Log-rank test with Bonferroni correction, p < 0.001) and +/– fish (p < 0.001). Mortality among the –/– individuals began at 30 dpf. At 90 dpf, only 20% of the original cohort were still alive. We note that when housed collectively, none of the –/– individuals survived to adulthood (Palmiotti et al. 2023). The improved survival (20%) of the –/– in this experiment might result from lack of competition for food with their more robust +/+ and +/– siblings.

Genotype also had a significant effect on fish growth from 24 to 44 dpf (**Fig. 2C**; Linear Mixed Model (LMM) length ∼ genotype + age, p < 0.001). The growth of the –/– fish was severely stunted: they were significantly shorter than the +/+ (Tukey’s HSD, p = 0.030) and +/– (p < 0.001) fish by 34 dpf and did not grow from 34 to 44 dpf (p = 0.235). By 90 dpf, the surviving –/– fish were over sixfold smaller by weight than the individuals of the other two genotypes (**Fig. 2D**; one-way ANOVA with Tukey’s HSD, p < 0.001). The –/– individuals were also paler, did not properly develop scales, and exhibited erratic swimming behavior (**Fig. 2E**, **Supplemental Movie S1**). Altogether, these results demonstrate that *bco1l* is essential for growth and survival during juvenile development in betta.

### bco1l knockouts are retinoic acid deficient

After documenting the time of death and altered morphology of –/– animals, we sought to characterize the biochemical consequences of the *bco1l* mutation. *bco1l* cleaves β-carotene into two molecules of retinal, which is then converted into retinoic acid (RA). Excess RA is degraded by *cyp26a1*, and the expression of this gene is boosted by RA (Hu et al. 2008; Isken et al. 2007; Samarut et al. 2015) making it a useful marker for relative RA concentrations (Isken et al. 2008). To test for evidence of altered RA levels in our *bco1l* mutants, we used digital PCR for absolute quantification of *cyp26a1* mRNA. There was no difference in *cyp26a1* expression between +/+, +/–, and –/– fish at 3 dpf, which is one day after hatching (**Fig. 3A**; one-way ANOVA, p = 0.156), suggesting that RA levels were equivalent among the genotypes and that *bco1l* is not required for RA production prior to 3 dpf. By 10 dpf, however, the –/– larvae had 6-fold lower *cyp26a1* than either the +/+ or +/– individuals (one-way ANOVA with Tukey’s HSD, p < 0.001), while +/+ and +/– did not differ in expression (p = 0.998). The same pattern was apparent at 20 dpf, with the –/– larvae having reduced *cyp26a1* expression relative to both the +/+ (p = 0.035) and +/– (p = 0.015) individuals. This pattern of expression across genotypes suggests that *bco1l* is required for RA production during larval, but not embryonic, development in betta.

**Figure 3.**
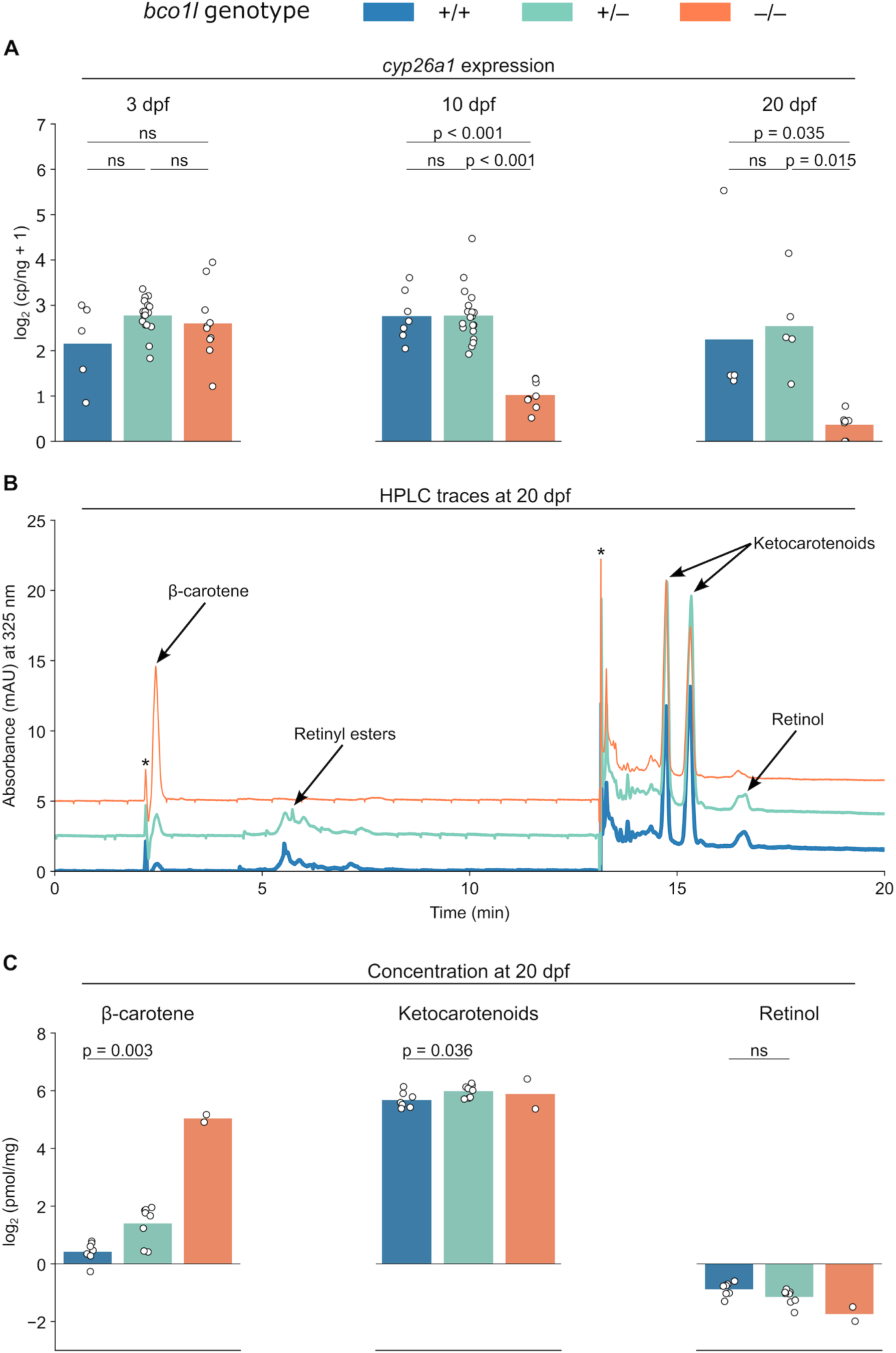
Retinoid and carotenoid content in embryonic and larval development in *bco1l* knockout betta fish. **A)** Whole-body *cyp26a1* expression at 3, 10, and 20 dpf across *bco1l* genotypes. p-values by one-way ANOVA with Tukey’s HSD. **B)** HPLC chromatograms across *bco1l* genotypes. Peaks corresponding to β-carotene, retinyl esters, ketocarotenoids, and retinol are indicated. * denotes peaks arising from a gradient solvent composition change. The +/– and –/– traces are vertically offset by 2.5 and 5 mAU, respectively. **C)** β-carotene, ketocarotenoid, and retinol concentrations in 20 dpf larvae of the three genotypes. p-values by Welch’s t-test.

To confirm that the reduced RA levels were due to the inability of the –/– larvae to convert β-carotene to VA and to compare ketocarotenoid concentrations between the genotypes, we quantified β-carotene, ketocarotenoids, and retinol in 20 dpf larvae of the three genotypes using high-performance liquid chromatography (HPLC) (**Fig. 3B, C**). The concentration of β-carotene was in +/– individuals was double that in +/+ (Welch’s two-sided *t*-test, p = 0.003). Furthermore, the –/– larvae displayed a mean concentration β-carotene 24 times higher than that of +/+ larvae. Ketocarotenoid content in +/– larvae was slightly higher than in +/+ larvae (p=0.036). There was no significant difference in retinol content between the +/+ and +/– groups (p = 0.063) whereas +/+ larvae had 1.8× more retinol than –/– larvae. Retinyl esters were not detected in the –/– larvae but were present in both the +/+ and +/– individuals (**Fig. 3B**). Altogether, the accumulation of β-carotene and the reduction of retinol and retinyl esters, indicate that *bco1l* is necessary for VA production in larval betta, explaining the deficit of RA in homozygous *bco1l* mutant animals.

### Retinoic acid rescues the bco1l mutation

Retinoic acid is a highly biologically active VA metabolite (Duester 2008; O’Byrne and Blaner 2013; Petkovich and Chambon 2022). To test for a causal relationship between the reduced RA levels in our knockouts and their stunted growth and death, we sought to rescue the –/– phenotype via exogenous RA supplementation. To that end, we housed larvae of the three genotypes in 5 nM RA (dissolved in a 0.1% DMSO solution) beginning at 5 dpf. Our chosen RA concentration is consistent with those of previous successful RA rescue experiments in developing zebrafish (Begemann et al. 2001; Wang et al. 2017). *bco1l* +/+ and +/– did not differ in their response to RA supplementation, so we combined these groups to increase our statistical power. In the control (DMSO only) group, expression of *cyp26a1* at 10 dpf in the +/+ plus +/– group was over 11-fold higher than in the –/– larvae (**Fig. 4A**; Welch’s t-test adjusted by Bonferroni correction, p < 0.001), confirming our prior observation (**Fig. 3A**) that –/– animals without RA supplementation have reduced RA levels. RA treatment increased *cyp26a1* expression in larvae of all three genotypes relative to their respective control groups (p < 0.001), demonstrating that exogenous RA was effectively absorbed by the larvae. Furthermore, when supplemented with RA, there was no significant difference in *cyp26a1* expression between the +/+ plus +/– group compared to –/– larvae (p = 0.343). This indicates that RA supplementation rescued the RA deficit and effectively equalized RA levels among fish with the three genotypes.

**Figure 4.**
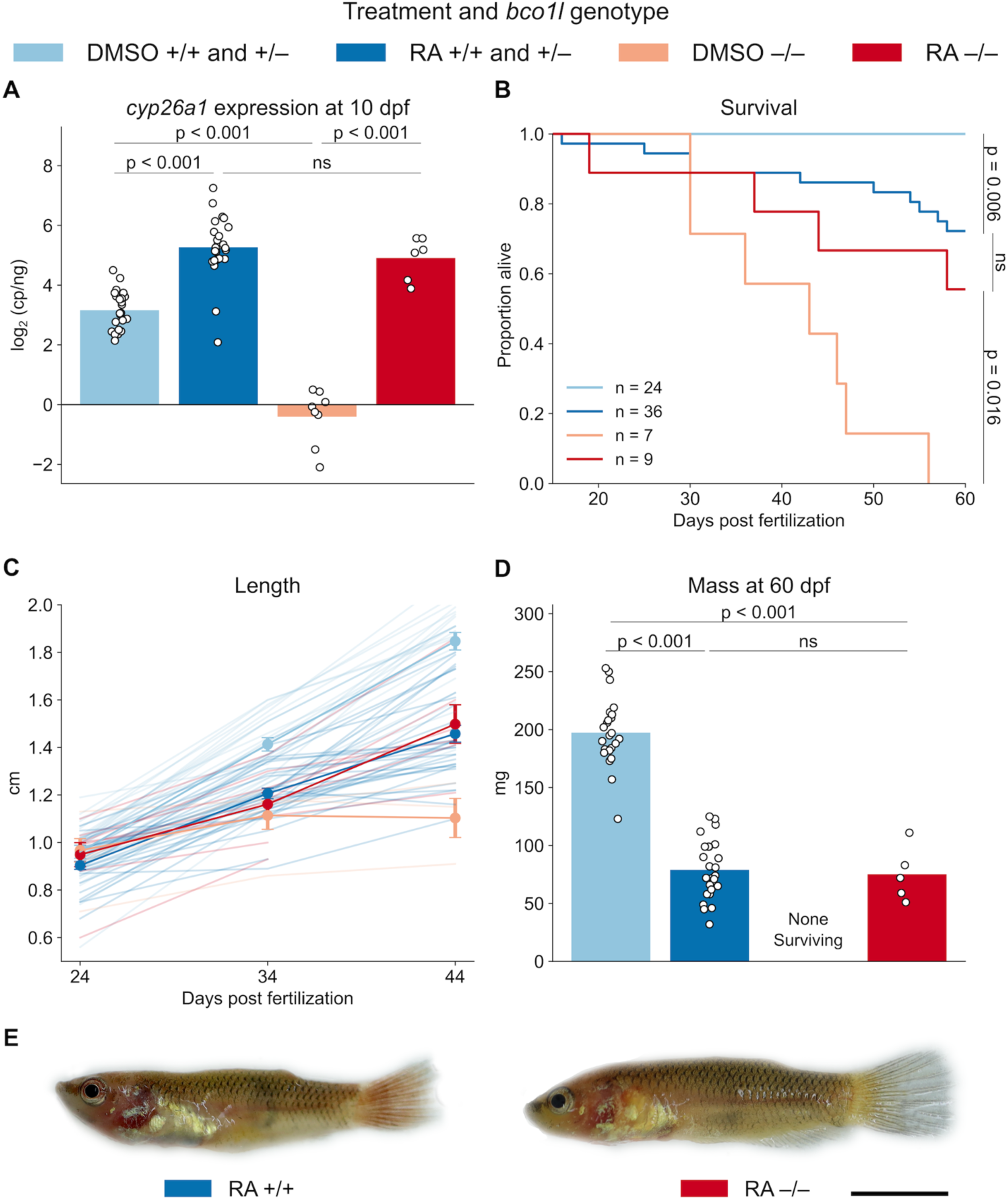
Exogenous retinoic acid supplementation rescues the *bco1l* knockout phenotype. **A)** Whole body *cyp26a1* expression in DMSO control and RA-treated cohorts of different genotypes at 10 dpf. p-values by Welch’s t-test adjusted by Bonferroni correction. **B)** Survival curves for DMSO control and RA-treated individuals from 15 to 60 dpf. p-values by Welch’s t-test adjusted by Bonferroni correction. **C)** Lengths of –/– and +/+ plus +/– fish in the DMSO control and RA-treated groups at 24, 34, and 44 dpf. Faint lines denote the growth of individual animals, circles denote the mean ± s.e.m. **D)** Masses at 60 dpf. p-values by Welch’s t-test adjusted by Bonferroni correction. **E)** Representative images of RA-treated +/+ and –/– animals at 60 dpf. Scale bar: 5 mm.

To assess the effect of RA treatment on growth and survival, we individually housed larvae in RA (dissolved in DMSO) or control (DMSO) from 15 to 60 dpf. RA significantly increased the survival of –/– fish (**Fig. 4B**; Log-rank test adjusted by Bonferroni correction, p = 0.016). Furthermore, there was no significant difference in survival on RA between the +/+ plus +/– group compared to –/– (p = 0.324), demonstrating that equalizing RA levels among genotypes equalized their survival. The +/+ plus +/– group on RA had increased mortality relative to their non-RA supplementation control (p = 0.006), which we attribute to toxicity of excessive RA (Wang et al. 2023; Chawla et al. 2018). Altogether, these results suggest that –/– fish die due to RA deficiency. RA supplementation also had a significant effect on growth (**Fig. 4C**; LMM length ∼ treatment/genotype group + age, p < 0.001). Control –/– individuals did not grow from 24 to 44 dpf (Tukey’s HSD, p = 0.445), while –/– on RA grew significantly (p < 0.001), demonstrating that RA treatment rescued growth. Furthermore, there was no significant difference in length between the +/+ plus +/– group on RA compared to –/– on RA at any time point (Tukey’s HSD, p > 0.05), and no difference in mass between these groups at 60 dpf (**Fig. 4D**; Welch’s t-test adjusted by Bonferroni correction, p = 0.754), indicating that RA eliminated growth differences between genotypes. We observed no apparent morphological differences among animals of the different genotypes treated with RA (**Fig. 4E**). Consistent with a toxic effect of RA, +/+ plus +/– animals treated with RA were significantly shorter than their control counterparts by 44 dpf (Tukey’s HSD, p < 0.001) and weighed significantly less at 60 dpf (Welch’s t-test adjusted by Bonferroni correction, p < 0.001). The results demonstrate that RA supplementation rescues both survival and growth of the *bco1l* mutation, indicating that the deleterious effects of the knockout are due primarily to RA deficiency. This is consistent with previous studies showing that insufficient VA during fish larval development can cause both stunted growth and death (Hernandez & Hardy, 2020).

### Expression of bco1 and bco1l and carotenoid metabolism during development

In addition to *bco1l*, *bco1* is also capable of catalyzing the conversion of β-carotene to retinal in ray-finned fish species (Lampert et al. 2003). To investigate why *bco1* does not compensate for the lack of *bco1l* in betta and to better understand how *bco1l* controls VA production, we characterized the expression levels and patterns of the two genes across development. Both *bco1* and *bco1l* were expressed in the intestine of betta larvae at 3, 5, and 10 dpf (**Fig. 5A**). At 3 dpf, *bco1* and *bco1l* expression overlapped in the intestinal epithelium. This pattern was consistently observed at 5 dpf, with both transcripts co-expressed in the developing intestine. By 10 dpf, co-expression of *bco1* and *bco1l* persisted in the intestine and was also detected in the liver. To measure the relative expression levels of the two genes in different tissues, we performed absolute quantification of *bco1* and *bco1l* transcripts using digital PCR in various tissues in adult betta, when animals are large enough to facilitate dissection of individual tissues. *bco1l* was 4-fold more abundant in the intestine than *bco1* (**Fig. 5B**; two-sided paired *t*-test, p = 0.006), consistent with the higher intestinal expression of *bco1l* observed in Atlantic salmon (Helgeland et al. 2014). *bco1* was 4-fold more abundant in skin (p = 0.017) and 2-fold more abundant in eye tissue (p = 0.009). The enriched skin expression is consistent with the effects of *bco1* mutation on skin pigment cells in zebrafish and pearl danio (Huang et al. 2021; Lampert et al. 2003). Both genes are expressed at similar levels in the gonads, head kidney, and liver. We did not detect expression of either paralog in muscle tissue, which contrasts with salmon, where both genes are expressed at low levels (Helgeland et al. 2014). Altogether, these results suggest that *bco1l* might be responsible for the bulk of β-carotene metabolism in the intestinal epithelium, where expression is detectable as early as 3 dpf and is higher than that of *bco1*in adults. By contrast, *bco1* activity may predominate in the eyes and skin.

**Figure 5.**
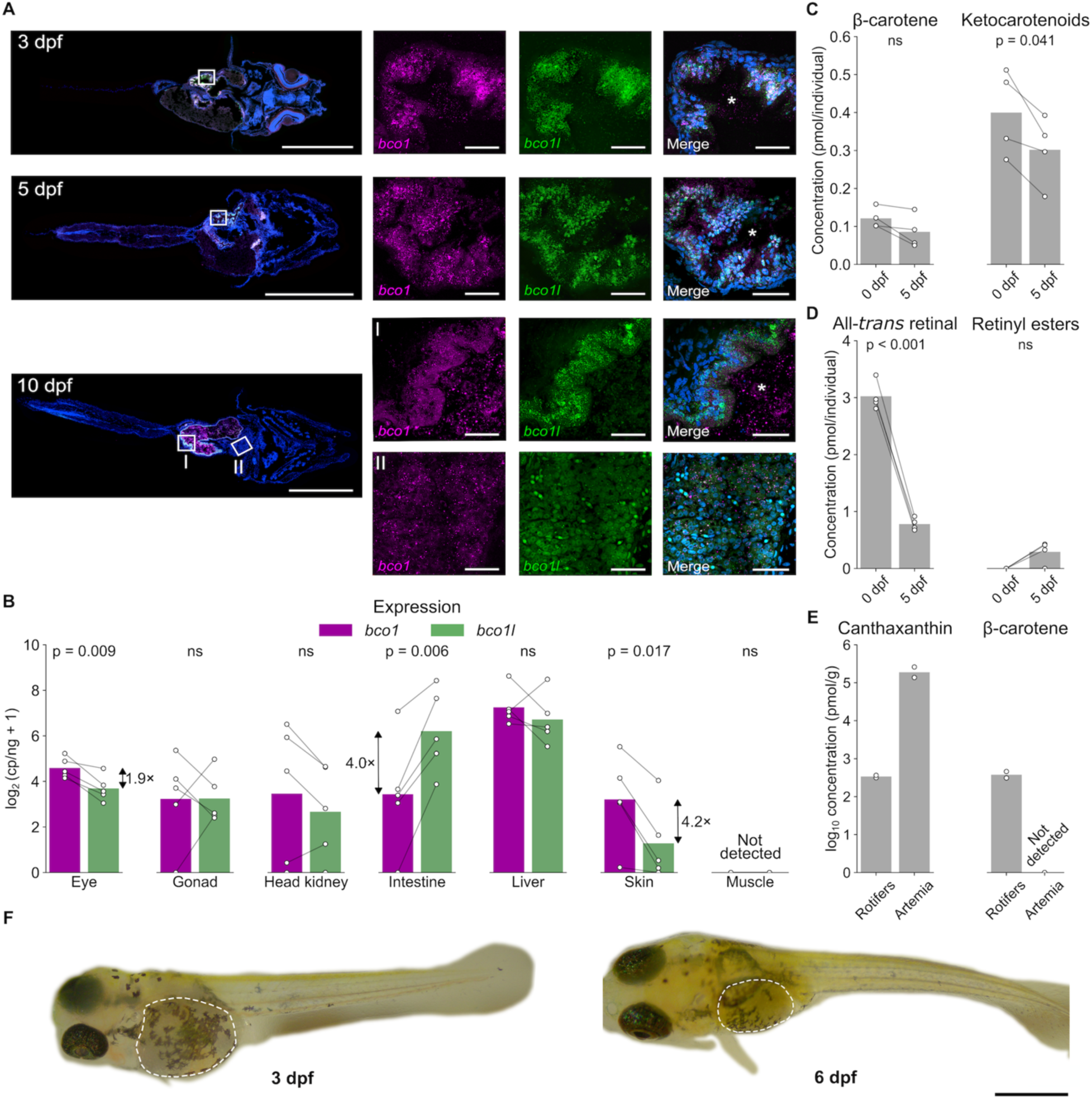
Spatiotemporal expression of *bco1* and *bco1l* and carotenoids and retinoids during embryonic development in betta fish. **A)** Detection of *bco1* and *bco1l* transcripts in wild-type larvae using *in situ* Hybridization Chain Reaction at 3, 5, and 10 dpf. **I.** Inset from 10 dpf highlighting the intestine. **II.** Inset from 10 dpf highlighting the liver. * denotes the intestinal lumen. Scale bars: 500 µm (whole larvae sections); 25 µm (magnified tissues). **B)** Expression of *bco1* and *bco1l* in various tissues in adult betta. p-values are by paired t-test. Data are paired by individual. **C, D)** Concentration of β-carotene and ketocarotenoids (**C**) and retinoids (**D**) in betta eggs and larvae at 5 dpf. p-values by paired t-test. Data are paired by cross. **E)** Concentrations of canthaxanthin and β-carotene in samples of rotifer and newly-hatched brine shrimp (*Artemia nauplii)* feeds. **F)** Representative images of 3 dpf and 6 dpf betta. Dashed lines indicate the yolk sac, which shrinks substantially as nutrients are consumed from 3 to 6 dpf. Scale bar: 1 mm.

RA is essential for pattern formation during embryonic development (Berenguer & Duester, 2022). We therefore sought to characterize the generation and utilization of VA (the RA precursor) through embryonic development in betta and to determine why lack of *bco1l* expression does not affect RA concentrations prior to 3 dpf but reduces them dramatically by 10 dpf (**Fig. 3A**). We used HPLC to separate polar and non-polar carotenoids and retinoids in eggs and 5 dpf larvae (before they start food intake) of developing betta (**Supplemental Fig. S2**). We also quantified carotenoids and retinoids in rotifers, which we feed betta fish from 5–14 dpf, and in newly-hatched brine shrimp (*Artemia nauplii*), which we feed betta from 15 dpf onwards. There was no significant difference in β-carotene concentration between eggs and 5 dpf larvae (**Fig. 5C**; two-sided paired *t*-test, p = 0.083) while total ketocarotenoid content was ∼25% lower in the 5 dpf larvae than the eggs (p = 0.041). To test whether betta embryos and larvae may use retinoids stored in the yolk, we also measured all-*trans* retinal and retinyl ester levels. The concentration of all-*trans* retinal in eggs was 7.5–25× higher than that of ketocarotenoids and β-carotene. By 5 dpf, the all-*trans* retinal concentration had decreased by 76% (**Fig. 5D**; p < 0.001). In zebrafish, the majority of yolk retinal is used up to support the development of the photoreceptors in the eyes (Isken et al. 2007; Lampert et al. 2003), a process likely conserved in betta. Retinyl esters were not detected in eggs, suggesting that betta, like other fish, store maternal VA as retinaldehyde bound to vitellogenin (Irie and Seki 2002; Lampert et al. 2003) (**Fig. 5D, Supplemental Fig. S2A**). Retinyl esters increased slightly by 5 dpf in some animals, indicating a portion of this retinaldehyde is converted to retinyl esters by that time (**Fig. 5D, Supplemental Fig. S2B**). The much lower levels of carotenoids compared to all-*trans* retinal in the eggs, coupled with a reduction in all-*trans* retinal through development to 5 dpf suggests that larvae do not use yolk carotenoids as a significant source for retinoid production by 5 dpf and instead use all-*trans* retinal stored in the yolk.

As vertebrates obtain pro-VA carotenoids and/or VA retinoids from food throughout life, we also determined the pro-VA carotenoid content of the rotifers and *Artemia* that we feed our fish. In rotifers, we tentatively identified a major chromatographic peak as β-carotene based on its retention time and spectral profile (**Supplemental Fig. S2E**). We did not detect β-carotene in *Artemia* (**Fig. 5E, Supplemental Fig. S2F**). Both rotifers and *Artemia* contained canthaxanthin and an additional unidentified ketocarotenoid. Additional peaks did not match the spectral properties of known carotenoids or retinoids. Our results indicate that betta larvae have access to dietary β-carotene and ketocarotenoids after 5 dpf (when they start eating rotifers). We suggest that the intestinal conversion of β-carotene and ketocarotenoids (likely converted into β-carotene first by other enzymes) to retinal by *bco1l* becomes critical after this point as they are exhausting their yolk stores of VA. This explains the reduced RA levels in the 10 dpf and 20 dpf *bco1l* knockouts. The lack of β-carotene in *Artemia* means that our betta do not have dietary access to it after 15 dpf, yet the VA-deficiency phenotype of the *bco1l* knockout becomes increasingly severe after this point (**Fig. 2**). As *Artemia* contain very little preformed VA (Moren et al. 2005), this indicates that betta derive VA from ketocarotenoids such as canthaxanthin, (a process demonstrated in other fish (Moren et al. 2002), and that *bco1l* is involved in this pathway. Using the same biochemical methods by which we demonstrated that *bco1l* cleaves β-carotene into two molecules of retinal (Kwon et al. 2022), we found no evidence of *bco1l* cleavage of canthaxanthin, suggesting that ketocarotenoids are first converted into β-carotene by other enzymes.

### Evolution and conservation of bco1 and bco1l

While one of a pair of paralogous genes is typically pseudogenized (Lynch and Conery 2000), *bco1* and *bco1l* are found across ray-finned fishes (Helgeland et al. 2014). Our identification of an essential function for *bco1l* in VA production and the differences in expression between *bco1* and *bco1l* prompted us to investigate the emergence of *bco1l* in the phylogenetic history of vertebrates.

*ninaB* is the invertebrate ortholog to *bco1* and *bco1l* and acts as a carotenoid oxygenase (von Lintig and Vogt 2000). It also acts as a retinoid isomerase, a function performed by *rpe65* in vertebrates (Oberhauser et al. 2008). We found *bco1* in all major vertebrate taxa, but *bco1l* was restricted to ray-finned fishes (**Fig. 6A**).

**Figure 6.**
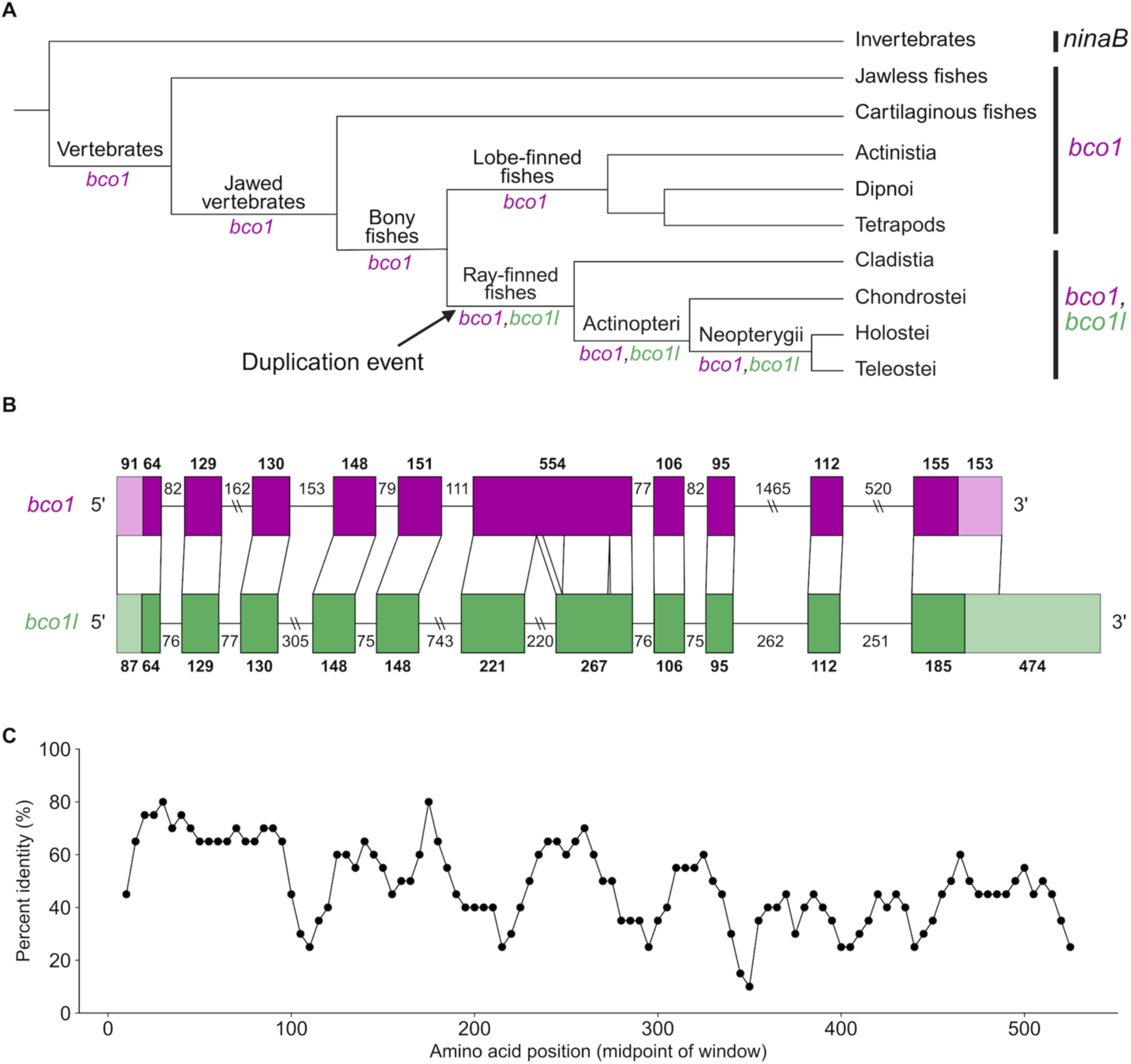
*bco1l* emerged from a duplication event in the stem lineage of ray-finned fishes. **A)** Vertebrate phylogeny showing presence of *bco1* and *bco1l* across lineages. *ninaB* is the invertebrate homolog. **B)** Exon-intron structure of betta *bco1* and *bco1l*. Lines connecting exons between the two genes indicate homologous regions. **C)** Percent identity of betta *bco1* and *bco1l* proteins. Points denote the percent identity of 20 amino acid windows with a 5 amino acid slide. The overall percent identity between the proteins is 47.7%.

Next, we probed the evolutionary origin of *bco1l* in more detail. If *bco1l* had arisen through a retrotransposition event, it would have no introns. However, the exon-intron structure of *bco1l* is conserved with *bco1* (**Fig. 6B)**. Furthermore, ancestral state reconstructions place *bco1* and *bco1l* adjacent to each other at the base of ray-finned fishes (see Methods). A parsimonious explanation for these observations, along with the phylogenetic pattern, is that *bco1l* emerged through a segmental duplication event in the stem lineage of the ray-finned fishes (**Fig. 6A**). Alignment of the amino acid sequences of betta *bco1* (547 residues) and *bco1l* (534 residues) revealed similarity across the lengths of the proteins, with an overall percent identity of 47.7% (**Fig. 6C**). This indicates that the duplication included the entirety of the ancestral *bco1* protein, and that no particular region has diverged more than others.

While both *bco1* and *bco1l* have been identified in numerous fish species, an inclusive analysis of their conservation across *Actinopterygii* taxa has not been conducted. To determine the evolutionary persistence of *bco1* and *bco1l*, we looked for evidence of each gene in 222 *Actinoptergii* species using publicly available reference genomes. To minimize false negatives, we excluded species for which one of the genes was not identified that had reference genomes with a BUSCO total copy percentage lower than 90%. We identified *bco1* orthologs in all analyzed species, and *bco1l* orthologs in 218 of 222 (**Supplemental Table S1**). We failed to identify *bco1l* in the following four species: *Etheostoma spectabile*, *Gymnodraco acuticeps*, *Notolabrus celidotus*, and *Pseudochaenichthys georgianus*. However, it is unclear whether this represents true evolutionary losses or errors in the reference genome assemblies (see Methods). Overall, our analysis demonstrates that both *bco1* and *bco1l* have been preserved throughout the ray-finned fishes.

## Discussion

### A model for vitamin A production via bco1l during development

By knocking out *bco1l* in betta fish, we discover that it is essential for VA and RA production. Lack of *bco1l* leads to stunted growth and death during development primarily due to insufficient RA. RA deficiency manifests after larvae have utilized the all-*trans* retinal stores in the yolk. At this stage, larvae transition to metabolism of dietary pro-VA carotenoids for continued retinoid biosynthesis and this metabolism requires *bco1l* (**Fig. 7**). Food sources rich in preformed VA may facilitate larval survival, yet the conservation of *bco1l* throughout ray-finned species suggests it has an essential role in nature.

**Figure 7.**
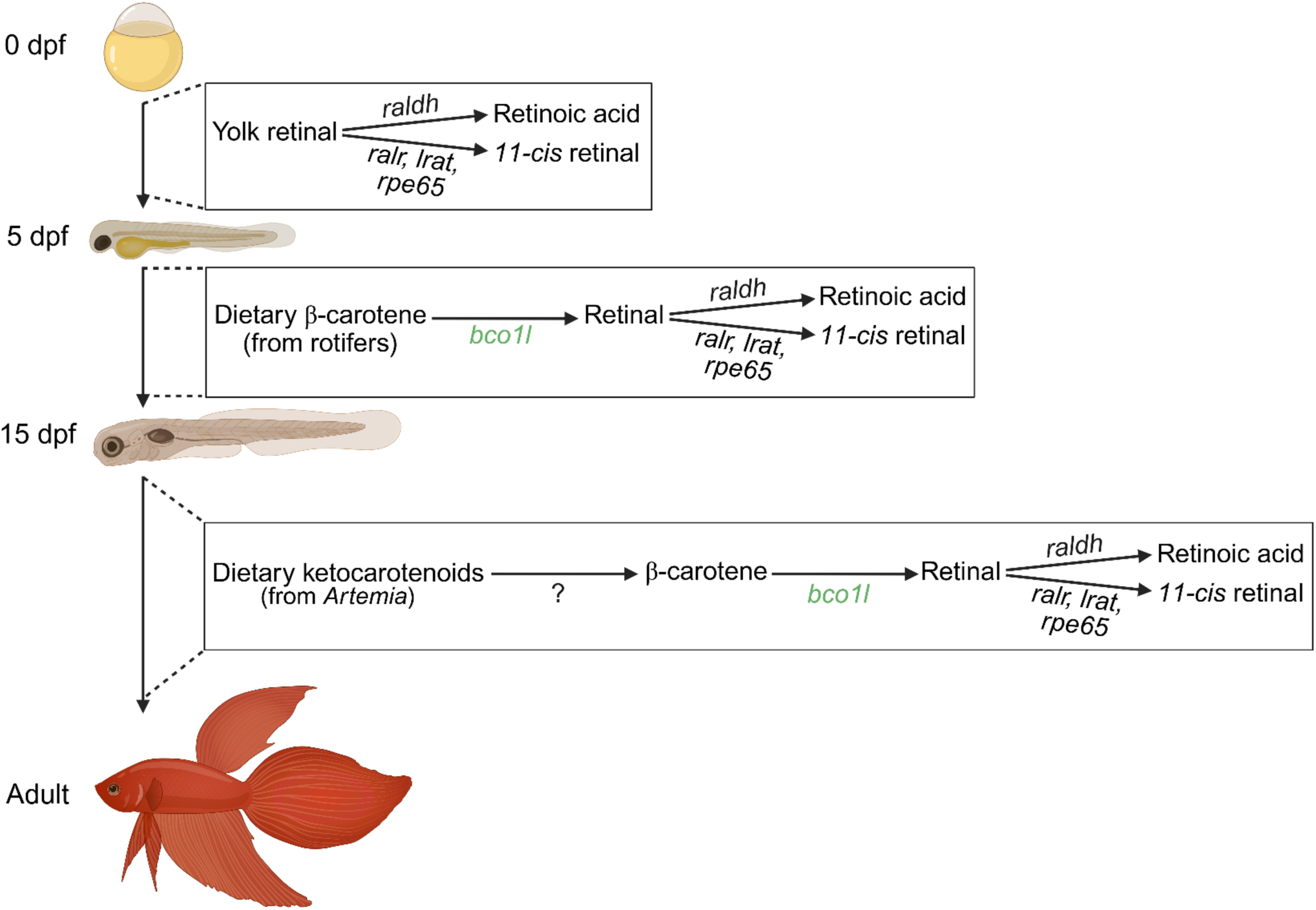
Vitamin A production during betta development. From 0 to 5 dpf, maternally deposited retinal in the yolk is utilized to produce the essential metabolite retinoic acid and the visual chromophore (*11-cis* retinal). Once the yolk stores of retinal are depleted, developing betta transition to retinoid production through the metabolism of dietary carotenoids. From 5-15 dpf, betta convert β-carotene from rotifer feeds to retinal via the 15,15’ oxygenase activity of *bco1l*. From 15 dpf to adulthood, betta rely on the provitamin A ketocarotenoid content of *Artemia* feeds. These molecules are likely converted to β-carotene through an unknown pathway and subsequently cleaved by *bco1l*. Created in BioRender.

In addition to β-carotene, ketocarotenoids including canthaxanthin are known to have pro-VA activity in fish, but the metabolic pathway of ketocarotenoid conversion to VA is unknown (Helgeland et al. 2019; Moren et al. 2002). Consistent with pro-VA activity of ketocarotenoids, wild-type betta showed no obvious signs of VA deficiency on a diet of *Artemia*, which contains canthaxanthin but no β-carotene nor preformed retinoids. By contrast, betta without *bco1l* are RA deficient and do not survive under this dietary condition. We therefore suggest that betta can convert ketocarotenoids to retinoids, in a process involving *bco1l* gene function. We found no biochemical evidence for ketocarotenoid cleavage by *bco1l*, in contrast to its ability to cleave β-carotene (Helgeland et al. 2019; Kwon et al. 2022). Therefore, ketocarotenoids are likely first converted to β-carotene and then cleaved to canonical retinoids, a pathway described in goldfish (Elshafey et al. 2023) and in VA-deficient rats (Sangeetha and Baskaran 2010).

### Mechanisms underlying the bco1l knockout phenotype

RA acts as a ligand for nuclear receptors (RARs), exerting transcriptional control over a variety of target genes. During embryonic development, a spatiotemporal gradient is established via the production and degradation of RA across different cell types, controlling anteroposterior patterning, limb initiation, heart development, and eye morphogenesis (Begemann et al. 2001; Berenguer et al. 2018; Berenguer and Duester 2022; Cunningham et al. 2013; Cunningham and Duester 2015; Grandel et al. 2002; Keegan et al. 2005; Sandell et al. 2007). As a result, disrupted RA signaling in embryonic zebrafish results in a curved body axis, malformed pectoral fins, cardiac edema, and microphthalmia (Begemann et al. 2001; D’Aniello et al. 2015; Isken et al. 2008; Lampert et al. 2003; Le et al. 2012). The absence of these defects in *bco1l* knockouts is consistent with the normal retinoid levels in wild-type and mutant betta during embryonic development. However, RA signaling is also important later in life, regulating apoptosis, osteoblast and osteoclast function, and immunity (Chawla et al. 2018; Green et al. 2016; Noy 2010; Raverdeau and Mills 2014). Our rescue experiments demonstrated that *bco1l* knockout phenotypes are caused by RA deficiency during larval and juvenile development, since treatment with exogenous RA improved survival.

### Deposition of carotenoids into eggs varies across fish species

Consistent with our results in betta, zebrafish eggs contain retinal, β-carotene, and various ketocarotenoids (Lampert et al. 2003). By contrast, Chinook salmon (*Oncorhynchus tshawytscha)* eggs contain little β-carotene and high concentrations of the ketocarotenoid astaxanthin (Li et al. 2005), which decline substantially during early development (Tyndale et al. 2008). The eggs of another salmonid, brown trout (*Salmo trutta*), contain relatively little astaxanthin and more lutein and zeaxanthin than *O. tshawytscha*, though still little to no β-carotene (Wilkins et al. 2017). Thus, pro-VA carotenoid deposition in egg yolk is rather variable. Differences in the amount and variety of carotenoids and retinoids available during embryonic development could influence RA production and the role of *bco1l* across species.

### Evolution and functions of bco1 and bco1l

Our phylogenetic analyses indicate that *bco1l* emerged via a gene duplication event in the stem lineage of the ray-finned fishes, updating prior interpretations of the origins of the *bco1* gene family (Helgeland et al. 2014). One of a pair of paralogous genes is usually psuedogenized shortly following duplication (Lynch and Conery 2000). This is particularly true in cases of small-scale duplication as it often disrupts the balance of interacting gene products, resulting in deleterious effects that drive the non-functionalization of one paralog (Birchler & Veitia, 2012; Wilson & Liberles, 2023). Nonetheless, we find both *bco1* and *bco1l* to be highly conserved across *Actinopterygii*, representing a useful case study for the evolution and preservation of duplicate genes.

Our *bco1l* knockout in betta provides insight into the relative function of *bco1* and *bco1l*. VA deficiency in *bco1l* mutant fish is not compensated by *bco1*, indicating that the two genes are not functionally redundant. In *Sarcopterygii* species, including humans, *bco1* converts the majority of dietary pro-VA carotenoids to retinoids in the intestinal mucosa (Lindqvist and Andersson 2002; Takitani et al. 2006). These compounds are then transported to the liver, from where they can be distributed through the blood to other tissues (Lindqvist and Andersson 2004; von Lintig 2010; von Lintig et al. 2005). While both *bco1* and *bco1l* are expressed in the intestine and liver of developing betta, *bco1l* is 4× more highly expressed in the intestine of adult fish than *bco1*, consistent with the expression pattern observed in Atlantic salmon (Helgeland et al. 2014). This indicates that *bco1l* may be primarily responsible for the intestinal metabolism of carotenoids in ray-finned fishes.

In addition to its role in the intestinal processing of carotenoids, mammalian *bco1* is expressed in various tissues such as the kidney, eye, and lungs, where it is thought to act in the local synthesis of retinoids (Lindqvist & Andersson, 2004; Paik et al., 2001; Yan et al., 2001). In zebrafish, *bco1* is expressed in a distally enriched gradient in the caudal fin (Rabinowitz et al. 2017), and morpholino mediated knockdown of the gene has cell-type specific effects (Lampert et al. 2003), providing further support for localized retinoid biosynthesis by *bco1*. Consistent with this, we observe *bco1* expression in various tissues in adult betta, with higher expression than *bco1l* in the skin and eyes. The tissue-specific synthesis of retinoids by *bco1* has been implicated in embryonic development in several species, including zebrafish (Kim et al. 2011; Lampert et al. 2003; Mora et al. 2004). A model for the subfunctionalization of these two genes relative to the ancestral *bco1* might therefore resemble the following: *bco1* acts to provide a local source of VA to tissues, such as those containing cells derived from the neural crest, while *bco1l* generates VA in bulk for storage and mobilization through its expression in the intestinal epithelium. This is consistent with the apparently normal embryonic development of betta *bco1l* knockouts and their manifestation of RA deficiency only following the switch to reliance on dietary carotenoids.

We note that the function of *bco1* in embryonic development in zebrafish (*Danio rerio*) has only been tested using morpholino-mediated knockdown, which have potential off-target effects (Lampert et al. 2003). In pearl danio (*Danio albolineatus*), a close relative of zebrafish, a *bco1* loss-of-function mutation only leads to a subtle fin-pigmentation phenotype in adults, with no reported deficits in viability or fertility (Huang et al. 2021). Further developmental and biochemical studies of *bco1* in betta and of *bco1* and *bco1l* in other ray-finned fishes will provide more details about their functions.

### Conclusions

Our findings suggest that the conversion of dietary pro-VA carotenoids to retinal via *bco1l* is essential for retinoic acid-dependent processes in larval and juvenile betta fish. Our results demonstrate that *bco1l* is not redundant with *bco1* and that it has been preserved in the genomes of ray-finned fishes since its emergence in their stem lineage.

## Materials and Methods

### Animal husbandry

Fish care followed our previously published protocols (Lichak et al. 2022), unless otherwise specified. Columbia University Institutional Animal Care and Use Committee approved all animal protocols. We began feeding larvae with *Brachionus rotundiformis* rotifers at 5 dpf. Rotifers were raised in reverse-osmosis (RO) water raised to a salinity of 15 ppt with Instant Ocean aquarium salt. We fed rotifers algae (Lichak et al. 2022). Each day, we decanted 4 L of a 200–600 rotifer/mL solution through a 41 µm sieve and resuspended rotifers in 250 mL of rotifer water in a 1 L beaker. We then mixed 25 mL aliquots of this concentrated solution with 175 mL fish water (Lichak et al. 2022). We fed this solution to our larval fish once per day (∼6.7 mL per fish). We transitioned betta to newly-hatched *Artemia* at 15 dpf. We used E-Z Egg brine shrimp eggs (Brine Shrimp Direct, Ogden, UT). We seeded egg batches daily by adding 45 mL of E-Z Eggs to 16 L of RO water with 2 cups Instant Ocean aquarium salt in a 16 L acrylic hatching cone. We allowed 18–24 hours for hatching prior to harvest, then drained the water through a 120 µm screen. We then resuspended hatched brine shrimp in 1 L of RO water. We stored brine shrimp at 4 °C for up to 24 hours (until the next harvest). Apart from experiments in which we individually housed larvae (see “Survey through development” and “Retinoic acid treatment”), we fed fish with *Artemia* twice per day (∼0.35 mL of the of the resuspended hatched brine shrimp per fish per feeding) from 15–29 dpf. From 30–79 dpf, we fed fish *Artemia* three times per day (∼0.35–1 mL per fish per feeding) and Golden Pearl Reef and Larval Fish pellets (Brine Shrimp Direct, Ogden, UT) once per day (100–300 µm; 3–9 mg per fish). From 80 dpf onwards, we fed adult fish 10 mL of *Artemia* and 3–5 Golden Pearl fish pellets (300–800 µm diameter) once per day.

### Genotyping

To extract DNA, we digested fin clips or whole embryo/larvae samples in 20 µL of a Proteinase K working buffer (10 mM Tris (pH 8), 2 mM EDTA, 0.2% Triton-X, 2 µg Proteinase K (D3001-2-H, Zymo Research)) at 65 °C for 40 min followed by 95 °C for 10 min. We used the lysate directly for genotyping. In cases where we also extracted RNA, we obtained DNA and RNA from tissue samples using the Quick-DNA/RNA Miniprep Kit (D7001, Zymo Research). Prior to all fin clip procedures, we anesthetized fish in a 0.8% tricaine methanesulfonate (MS-222) solution in fish water.

For wild-type +/+ × heterozygous +/– outcrosses, we used a T7 endonuclease assay (Mashal et al. 1995; Sentmanat et al. 2018) (M0302, New England Biolabs). We amplified *bco1l* from genomic DNA using the following primers: CAGATCCCTGCCAGAACATT and TGAGCCTGTGGTTCACTGAC. We induced hybridization by running 10 µL unpurified PCR product with 7.75 µl water and 2 µl NEB Buffer 2 (B6002, New England Biolabs) in the following conditions: 98 °C (5 min), 95–85 °C (–2 °C/sec), 85–25 °C (–0.1 °C/sec). We then added 0.25 µl T7 Endonuclease I directly to each of the hybridized samples and incubated at 37 °C for 60 min. T7 endonuclease I cuts heteroduplex DNA (arising from heterozygosity), which we visualized after electrophoresis in a 1.5% agarose gel.

For +/– × +/– crosses, we genotyped offspring using a digital PCR assay with primers specific to *bco1l* (CCGCTATGGTGACGACTACTAC and TGCCGACAGTTTCCAAGGT) and probes specific to the wild and mutant alleles (Wild: /56-FAM/TCCTCTGAGATCAAC/3MGB-NFQ/; Mutant: /5SUN/CTCCTCTGATGATCAAC/3MGB-NFQ/). We performed all reactions using a QIAcuity digital PCR system (QIAGEN) and 8.5 k partition dPCR plates. We used either a TaqMan probe mix (Thermo Fisher Scientific) or reconstituted probes and primers from Integrated DNA Technologies (IDT). For the TaqMan probes, we used a 14 µL reaction volume with the following reagent concentrations: 3.75 µL 40× QIAcuity Probe Mastermix (QIAGEN, 250102), 0.375 µL 40× TaqMan assay mix, 8.875 µL RNase/DNase free water, and 1 µL template DNA (6 ng/µL for extracted DNA or 1 µL of product solution from the Proteinase K digestion). For the IDT probes and primers, we used a 14 µL reaction volume with the following reagent concentrations: 3.75 µL 40× QIAcuity Probe MM, 0.135 µL *bco1l* forward primer (100 µM), 0.135 µL *bco1l* reverse primer (100 µM), 0.03 µL *bco1l* wild-type probe (100 µM), 0.03 µL *bco1l* mutant probe (100 µM), 8.92 µL RNase/DNase free water, and 1 µL template DNA. We used the QIAGEN Probe Priming protocol and the following cycling conditions: 95 °C (2 min), 40×[95 °C (15 s), 60 °C (30 s)]. We performed fluorescence imaging with the green (FAM; wild) and yellow (VIC; mutant) channels with an exposure duration of 500 ms and a gain of 6. We set fluorescence thresholds to 30 Relative Fluorescence Units (RFUs). We included +/+, +/–, and –/– control samples in every run.

### Survey through development

We sampled individuals from a +/– × +/– cross at 3, 6, 10, 12, and 20 dpf using a glass Pasteur pipette cut at the taper and flame-smoothed. We imaged each fish using a Leica S9i Stereo Microscope and then genotyped each individual. All images included a ruler to compare length and were scrutinized for evidence of abnormal phenotypes.

At 22–24 dpf, we individually housed larvae of all three genotypes in 200–300 mL of fish water in opaque plastic cups to prevent death due to competition for food. We fed the fish *Artemia* 4× per day (∼20–50 shrimp per feeding) and replaced the media every other day. Twice per day, we screened for dead fish. If a fish was found dead, we stored it at –20 °C for genotyping and recorded its age at death. We photographed each fish in the cups top-down at 24, 29, 34, 39, and 44 dpf. We processed the images using ImageJ (Schneider et al. 2012), obtaining length measurements by multiplying the ratio of the number of pixels along the body of the fish to those along the diameter of the bottom of the cup by the known diameter of the cup bottom (4.5 cm). At 90 dpf, we weighed and genotyped all remaining fish. Fish were euthanized via submersion in ice water, patted dry using a Kimwipe, and weighed using a Mettler-Toledo ML104T/00 Analytical Balance. We took all photos and videos using a Canon EOS RP camera with a Canon Macro lens EF 100 mm inside a photo tent illuminated with white light-emitting diodes.

### Retinoic acid treatment

We dissolved all-*trans-*Retinoic acid (RA; Sigma-Aldrich R2625) in dimethyl sulfoxide (DMSO) at a concentration of 0.1 M and stored it in 5 µL aliquots in the dark at –70 °C. Beginning at 5 dpf, we housed treatment larvae in fish water containing 5 nM RA and 0.1% DMSO. Control larvae were housed in 0.1% DMSO. At 10 dpf, we sampled treated individuals and quantified *cyp26a1* expression across the three genotypes. To prevent competition for food, we individually housed both the treatment and control fish in plastic cups at 15 dpf containing 200–300 mL fish water. We fed fish with *Artemia* 2× per day (∼20-50 fish per feeding) and replaced the media every other day. We tracked survival, imaged each fish at 24, 34, and 44 dpf, and obtained length measurements using the same protocol as for the untreated fish (see “Survey through development”). We terminated the experiment at 60 dpf and genotyped and weighed the remaining fish.

### Digital PCR for cyp26a1, bco1, and bco1l

We extracted total RNA from whole body samples of embryonic and larval fish at 3, 10, and 20 dpf using the Quick-DNA/RNA Miniprep Kit (D7001, Zymo Research). We extracted total RNA from eye, gonad, head kidney, intestine, liver, skin, and muscle samples of five adult (3 male, 2 female), wild-type betta of the red veil variety using the Quick RNA Miniprep Kit (R1054, Zymo Research). We quantified RNA using a Nanodrop One (13-400-519, Thermo Fisher Scientific, Waltham MA) and a Qubit RNA IQ Assay (Q33216, Invitrogen Qubit 3.0 Fluorometer). We synthesized cDNA using the LunaScript RT SuperMix kit (E3010, New England Biolabs).

In embryo and larval samples, we quantified *cyp26a1* expression using a PrimeTime qPCR Probe Assay (IDT) with primers (GCTGGTGGAGGCTTTTGAGG, GTGTCGGACTCCTGCACCTT) and probe (/56-FAM/TGTACAGGG/ZEN/GTCTGAAGGCGAGGAA/3IABkFQ/) specific to the mRNA transcript of *cyp26a1*. We quantified *bco1* and *bco1l* expression in adult tissue samples using two PrimeTime qPCR Probe Assays (IDT) with primers (*bco1*: TGACACAGAAACCAAGGAACT, GACCGCGCCATCATCTT; *bco1l*: GAGAACTGCTACCCATCAGAAC, GGAAATGTGAGGATCTGGAGAAA) and probes (*bco1*: /5TexRd-XN/AGACAACTGCTTTCCATCGGAACCA/3BHQ_2/; *bco1l*: /56-FAM/TCACACCAT/ZEN/CATCCTCATCCACAGC/3IABkFQ/) mRNA transcripts of *bco1* and *bco1l*.

We performed all reactions using a QIAcuity digital PCR system and 8.5 k partition dPCR plates. For *cyp26a1*, each reaction contained 50 ng of RNA reversed transcribed into cDNA in a 15 µL of solution (0.75 µL 20× qPCR probe mix, 3.75 µL QIAcuity Probe Mastermix, 5 µL 10 ng/µL cDNA template, 5.5 µL RNase/DNase free water). For *bco1* and *bco1l*, we ran multiplexed dPCRs with 40.2 ng RNA/cDNA input in 15 µL of solution (0.75 µL 20× *bco1* qPCR probe mix, 0.75 µL 20× *bco1l* qPCR probe mix, 3.75 µL QIAcuity Probe Mastermix, 6.7 µL 6 ng/µL cDNA template, 3.05 µL RNase/DNase free water). For both assays, we used the QIAGEN Probe Priming protocol and the following cycling conditions: 95 °C (2 min), 40×[95 °C (15 s), 60 °C (30 s)]. We performed fluorescence imaging using the green (FAM; *cyp26a1, bco1l*) and red (TexRd; *bco1*) channels with an exposure duration of 500 ms and a gain of 6. We set fluorescence thresholds to 40 RFU for *cyp26a1*, 100 RFU for *bco1,* and 80 RFU for *bco1l*.

### HPLC for carotenoid and retinoid concentrations

We collected between 50 and 100 eggs in indirect light within five hours of mating from each of four wild-type betta crosses of the red veil variety. We raised the remaining offspring in low light until 5 dpf, then sampled between 50 and 100 larvae from the same crosses. All samples were counted, pooled, and immediately stored in –70 °C in the dark. We sampled larvae of the *bco1l* knockouts at 20 dpf and genotyped them via a small caudal fin biopsy. We then made pools of 3–5 fish of the same genotype. We euthanized larvae via submersion in ice and patted dry with Kimwipes prior to weighing. We weighed samples using a Mettler-Toledo ML104T/00 Analytical Balance. All pooled samples were between 14–26 mg and stored at –70 °C in the dark. We sampled newly-hatched brine shrimp (*Artemia nauplii*) and rotifers from our lab cultures, concentrating them into a pellet in a microcentrifuge tube via centrifugation at 20,817 rcf. We removed excess water via a pipette after centrifugation. Samples were between 12 and 25 mg.

We homogenized whole larvae, eggs, brine shrimp, and rotifers using a handheld homogenizer in 100 μL of phosphate-buffered saline (PBS, pH 7.4). We extracted carotenoids and retinoids after the addition of 200 μL 2 M hydroxylamine (pH 6.8) and 200 μL methanol to form the corresponding retinaldehyde oximes (syn and anti). Then, we added 400 μL acetone, followed by sequential extraction by two rounds of 500 μL hexane. We separated the organic phase by centrifugation, collected and vacuum-dried using an Eppendorf vacuum centrifuge.

We reconstituted the debris in a 99:1 (v/v) mixture of hexane and ethyl acetate and subjected it to gradient elution via high-performance liquid chromatography (HPLC). We performed gradient separation in two steps: the first utilized a 99:1 (v/v) hexane:ethyl acetate mixture for 10 minutes, allowing the elution of β-carotene and retinyl esters. The second step employed an 80:20 (v/v) hexane:ethyl acetate mixture for 15 min, facilitating the elution of more polar compounds, including keto-carotenoids, retinal-oximes and retinol. We conducted HPLC analysis using an Agilent 1260 Infinity Quaternary HPLC system equipped with a quaternary pump (G1312C), integrated degasser (G1322A), thermostatted column compartment (G1316A), autosampler (G1329B), diode-array detector (G1315D), and ChemStation software for data acquisition and analysis. We carried out chromatographic separation at 25 °C using a Pinnacle DB Silica column (5 μm, 4.6 × 250 mm; Restek) at a constant flow rate of 1.4 mL/min. For quantification, we calibrated the HPLC system with authentic standards of known carotenoids and retinoids. Peak areas were integrated to determine the concentration of each compound in the samples. We purchased all-*trans* retinol and retinal standards from Toronto Biochemicals. Carotenoid standards, including β-carotene, echinenone, canthaxanthin, and astaxanthin were a gift from Dr. Adrian Wyss (DSM Ltd., Sisslen, Switzerland).

### HCR for bco1 and bco1l expression

We performed *in situ* hybridization chain reaction (HCR) to detect *bco1* and *bco1l* mRNA transcripts in wild-type betta larvae at 3, 5, and 10 dpf, using the standard protocol and reagents from Molecular Instruments (v3.0 protocol, Choi et al. 2018). We designed HCR probe sets following the method and code presented in (Kuehn et al. 2022), and probe sequences are available in **Supplemental Table S2**. We ordered probes as oPools Oligo Pools (IDT). We anesthetized fish in 0.8% MS-222 and euthanized by immersion in ice-cold water. We fixed samples overnight at 4 °C in 4% paraformaldehyde (PFA), then washed them three times in PBS. Samples were cryoprotected by stepwise incubation in 15% and 30% sucrose in PBS until they sank. We then embedded them in “optimal cutting temperature” (OCT) and stored at −70 °C until use. We sectioned each sample at 7 μm thickness in the horizontal plane using a cryostat. We post-fixed the slides in 4% PFA for 15 minutes at 4 °C. Our remaining steps followed the protocol described in the HCR™ Gold RNA-FISH user guide. We included negative controls by processing samples identically but omitting the probe during hybridization. We imaged the samples using a W1-Yokogawa Spinning Disk Confocal microscope. We applied linear adjustments to brightness and contrast uniformly to entire images.

### Conservation and evolution of bco1 and bco1l

We determined that *bco1l* arose by duplication of *bco1* in the stem lineage of the ray-finned fishes (*Actinopterygii*) based on the following: (1) *bco1* is the most similar paralog of *bco1l* according to Ensembl release 114 (May 2025) Gene Tree (Dyer et al. 2025); (2) *bco1l* and *bco1* are adjacent to each other in a head to tail orientation in the genomes of several ray-finned fishes that are not monophyletic, and Genomicus v110.01 (Nguyen et al. 2022) places these two genes adjacent to each other in its reconstruction of the ancestral state of ray-finned fishes; and (3) according to Ensembl, Genomicus, and NCBI Orthologs (Sayers et al. 2024), *bco1l* has orthologues in fish species from all *Actinopterygii* lineages (Cladistia, Chondrostei, Holostei, and Teleostei) but not in any non-*Actinopterygii* species; and (4) according to Ensembl, Genomicus, and NCBI Orthologs, *bco1* is present throughout vertebrates.

To analyze the exon-intron structure of *bco1* (transcript variant X1) and *bco1l* (transcript variant X1), we obtained the lengths of the exons and introns for each gene from NCBI. We then aligned the mRNA transcripts from both genes using Clustal Omega (Ahmad et al. 2025) and determined homologous regions for each exon. We annotated indels in either gene if they exceeded 8 nucleotides in length. We then aligned the protein sequences of both genes using Clustal Omega and determined the percent identity along the lengths of the proteins in 20 amino acid windows with a step size of 5 amino acids.

### Presence of bco1 and bco1l across Actinopterygii

We obtained the list of all 222 unique ray-finned species with an annotated reference genome with a BUSCO total copy percentage > 90% from NCBI (Sayers et al. 2024). We then checked whether NCBI had identified putative orthologs for *bco1* and/or *bco1l* for every species in the list. If an ortholog was identified, we listed that species as having that gene. In the remaining species for which NCBI did not identify and ortholog for one or both genes, we searched for the missing ortholog using the following steps: (1) we checked the Ensembl comparative genomics (release 113) database (Dyer et al. 2025) by searching for the gene name (*bco1* or *bco1l*) and the species or by locating the Ensembl gene in the NCBI genome data viewer; (2) we manually searched for the missing orthologs in the species’ NCBI reference genome using BLASTp and reciprocal best match with the betta *bco1* and *bco1l* amino acid sequences (Ward and Moreno-Hagelsieb 2014); (3) we checked for orthologs using BLASTp in any alternative annotated reference genomes for those species available from NCBI.

For species in which we failed to identify orthologs for one of the two genes using these methods, we blasted the *bco1* or *bco1l* nucleotide sequence (discontiguous megablast) of the most closely related species included in our analysis against any unannotated alternative reference genomes available from NCBI. If we found a significant hit, we individually blasted the nucleotide sequence of each exon in the gene of the closely related species against the species being analyzed and constructed the mRNA sequence of the missing gene. We then established an open reading frame by aligning the two mRNA sequences (that of the species being analyzed and their close relative) using Clustal Omega. We translated the mRNA sequence and checked for premature stop codons. Finally, we blasted the amino acid sequence of the newly annotated gene against the betta genome using BLASTp to ensure that the top hit was the expected gene. We identified a *bco1* ortholog for *Poecilia mexicana*, *Poecilia latipinna*, and *Poecilia formosa* as well as a *bco1l* ortholog for *Salarias fasciatus* using this method.

We failed to identify an ortholog for *bco1l* in *Notolabrus celidotus*, *Gymnodraco acuticeps, Etheostoma spectabile*, and *Pseudochaenichthys georgianus.* For these species, we blasted the nucleotide sequence of genes from two closely related species in the genomic vicinity of *bco1l* to check for reference genome integrity in the syntenic region. However, we were unable to navigate to the syntenic region in any of these species, making it difficult to determine whether the lack of *bco1l* represented a true evolutionary loss or an incomplete genome assembly.

### Statistical analyses

We performed all statistical tests in JMP Pro 18 or Rstudio v.2024.12.1 (R v.4.4.3) (R Core Team 2025). When we log-transformed data for visualization, we performed statistical analyses on the transformed datasets.

## Acknowledgements

This work was supported by the following grants: National Institutes of Health (NIH) R35GM143051 to A.B. and NIH grants EY020551 and EY028121 to J.v.L. Pei-Yin Shih provided technical assistance. Sam Szalkowski and Sebastian Lumapas helped with fish care and experiments.

## Author Contributions

L.S.K., A.P., J.v.L, and A.B conceived the study. L.S.K., P.R.V., S.B., and Y.Z., performed experiments. L.S.K and P.R.V analyzed data and produced figures. L.S.K, P.R.V, and A.B. wrote the paper with revisions from all authors. J.v.L. and A.B. provided funding and resources.

